## Supplementary figures and images for "Hexanematic crossover in epithelial monolayers depends on cell adhesion and cell density"

### Nematic and hexatic order depend on the cell line and monolayer density.

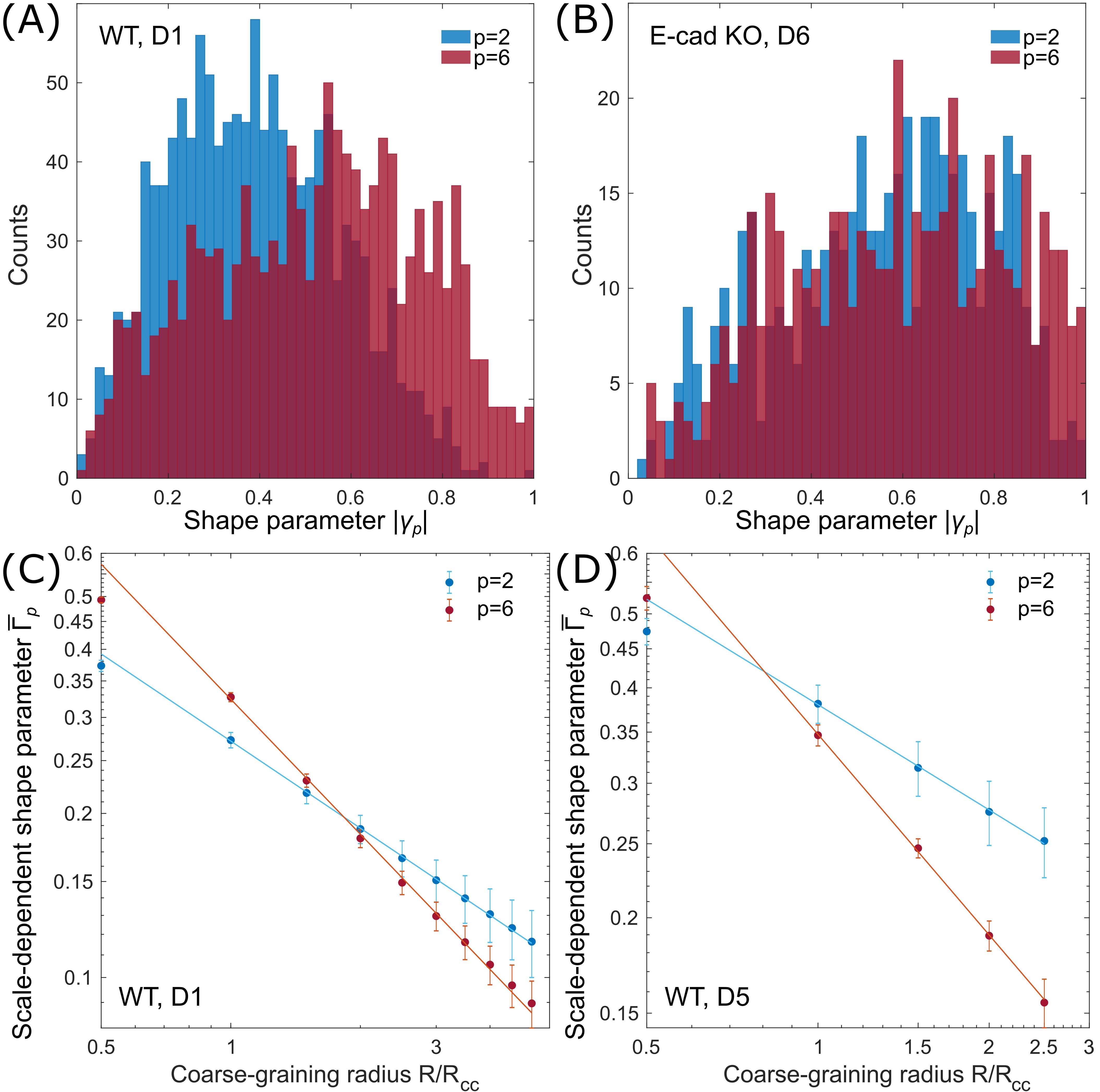

### The defect density depends on the coarse-grained orientation field.

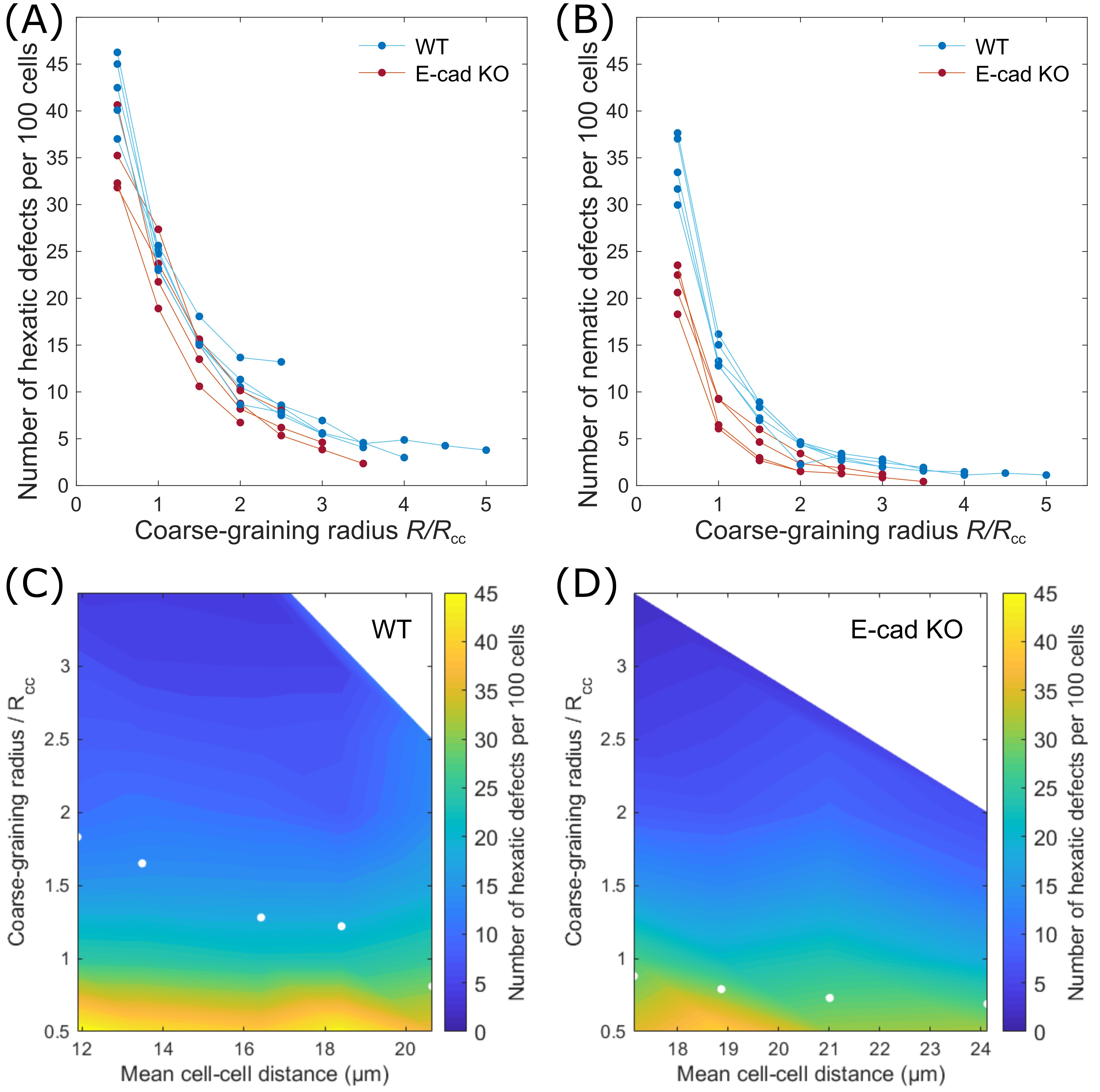

### The mean cell-cell distance increases with increasing substrate stiffness.

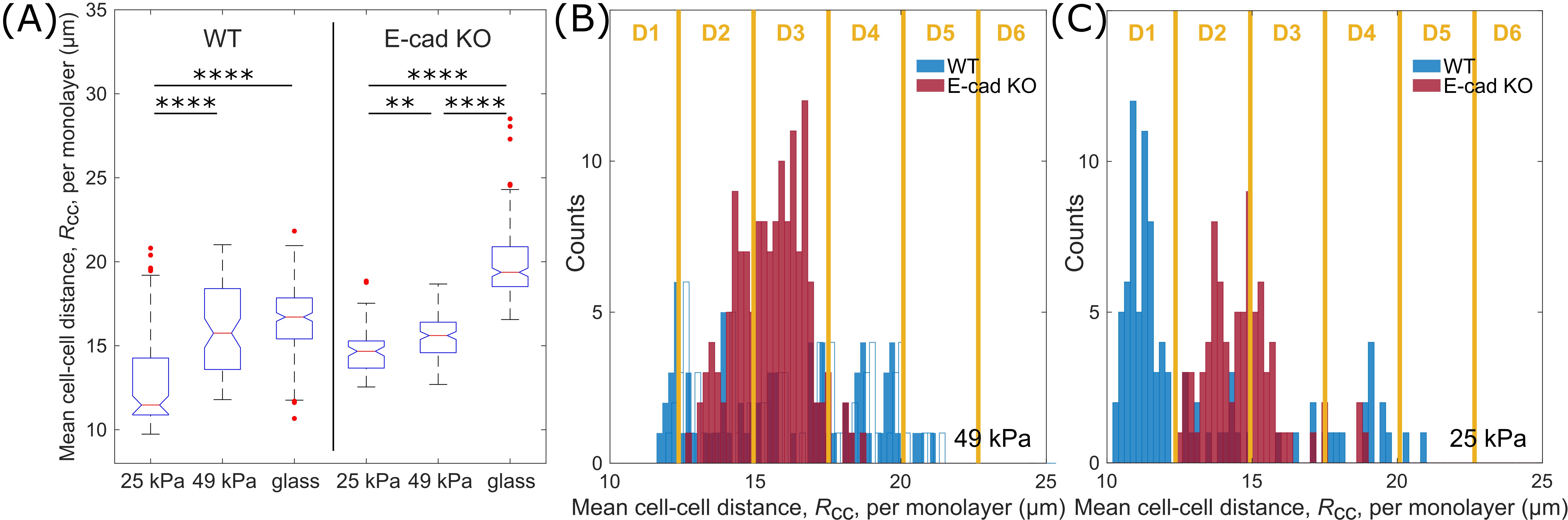
